# The hunger games: sensing host arginine is essential for *Leishmania* parasite virulence

**DOI:** 10.1101/751610

**Authors:** Adele Goldman-Pinkovich, Sriram Kannan, Roni Nitzan-Koren, Madhu Puri, Yael Bar-Avraham, Jacqueline A. McDonald, Aakash Sur, Wen-Wei Zhang, Greg Matlashewski, Rentala Madhubala, Shulamit Michaeli, Peter J. Myler, Dan Zilberstein

**Affiliations:** Faculty of Biology, Technion - Israel Institute of Technology, Haifa, Israel; The Mina and Everard Goodman Faculty of Life Sciences and Advanced Materials and Nanotechnology Institute, Bar-Ilan University, Ramat-Gan Israel; School of Life Sciences, Jawaharlal Nehru University, New Delhi, India; Center for Global Infectious Disease Research, Seattle Children’s Research Institute, USA; Department of Biomedical Informatics and Medical Education, University of Washington, Seattle, USA; Department of Microbiology and Immunology, McGill University, Montreal, Canada; Department of Global Health, University of Washington, Seattle, USA

**Author notes:** Department of Immunology and infectious Diseases, Harvard T.H. Chan School of Public Health, Boston, USA.

## Abstract

Arginine homeostasis in lysosomes is critical for growth and metabolism of mammalian cells. They employ a specific sensor (SLC38A9) that monitors intra-lysosome arginine sufficiency and subsequently up-regulates cellular mTORC1 activity. Lysosomes of macrophages (phagolysosomes) are the niche where the parasitic protozoan *Leishmania* resides and causes important human disease. Several years ago, we discovered that upon arginine starvation, cultured *Leishmania* parasites promptly activate a MAPK2-mediated Arginine Deprivation Response (ADR) pathway, resulting in up-regulation of the *Leishmania* arginine transporter (AAP3), as well as a small group of other transporters. Significantly, ADR is also activated during macrophage infection, implying that the intracellular parasite actively depletes arginine within the host phagolysosome, likely to prevent mTORC1 activation and enhance intracellular development. We hypothesize that ADR-mediated up-regulation of AAP3 activity is necessary to withstand the resultant arginine starvation. Both copies of the *AAP3* genes are located (in tandem) on a tetrasomic chromosome (chr31), but only one (*AAP3.2*) is responsive to arginine deprivation. CRISPR/Cas9-mediated disruption of the *AAP3* locus yielded mutants that retain a basal level of arginine transport (mediated by *AAP3.1*), but lack a functional copy of *AAP3.2* and are therefore not responsive to arginine starvation. While these mutants grow normally in culture as promastigotes, they were impaired in their ability to develop inside THP1 macrophages grown under physiological concentrations of arginine (0.1 mM). However, flooding the macrophage growth medium with arginine (1.5 mM) restored parasite infectivity and intracellular growth to that of wild type. The results indicate that inside the host macrophage, *Leishmania* must overcome the arginine “Hunger Games” by up-regulating transport of arginine *via* the ADR. Furthermore, the *AAP3.2* mutants were ~70-80% less virulent in Balb/C mice, showing, for the first time, that the ability to monitor and respond to changes in host metabolite levels is essential for pathogenesis.

## Introduction

The lysosomes of phagocytic cells are where pathogens fight for life in a hostile environment, dependent on the host for essential nutrient supply. In the case of the obligatory intracellular parasite *Leishmania*, arginine availability is of particular interest because it is important for both host defense and parasite proliferation. Sensing of arginine levels in the lysosome lumen is a key mechanism that regulates mTORC1 activity in mammalian cells (Saxton and Sabatini, 2017). In macrophages, mTORC1 activation induces a Th1 response, resulting in production of cytotoxic nitric oxide from arginine (Weinstein et al., 2000). This pathway is the major means by which macrophages kill invading pathogenic microorganisms. Interestingly, *Leishmania* parasites are able to counteract this outcome by activating a Th2 response, directing arginine towards polyamine biosynthesis instead of NO production, thereby enabling the parasites to persist and cause long-term non-healing infections (Shahi et al., 2013)(Rath et al., 2014). However, both Th1 and Th2 responses result in arginine depletion in the macrophage phagolysosome, presenting an existential threat to their survival. In this study, we explore the mechanism that enables intracellular *Leishmania* to win this “Hunger Game”.

Recently, our laboratory discovered that an MAPK2-mediated Arginine Deprivation Response (ADR) pathway is triggered immediately upon arginine starvation of the cultured extracellular insect-form (promastigote) *Leishmania*, resulting in up-regulation of arginine transporter (AAP3) mRNA and protein levels, with a concomitant increase in arginine transport efficiency (Goldman-Pinkovich et al., 2016). Significantly, the ADR is also activated during *Leishmania* infection of macrophages (Goldman-Pinkovich et al., 2016),(Pawar et al., 2019), suggesting a role for intra-lysosomal arginine sensing in intracellular (amastigote) parasite survival and pathogenesis. In this study, we sought to critically assess this hypothesis by creating mutants that are unable to up-regulate *AAP3* expression after arginine starvation.

## Results and discussion

Two genes (AAP3.1 and AAP3.2) that are highly conserved (only three amino acid differences) within their coding sequence (CDS) and 5’ untranslated region (UTR) but have very different 3’ UTRs encode AAP3. However, only *AAP3.2* is responsive to arginine deprivation (Darlyuk et al., 2009). Chromosome 31, where the two *AAP3* genes occur in a tandem array, is tetraploid in *L. donovani* (Downing et al., 2011); making classical homologous gene replacement approaches cumbersome and tedious, especially since the plasticity of *Leishmania* genome enables parasites to retain an additional wild-type chromosome as well as those containing the selectable marker(s)(Laffitte et al., 2016). Therefore, we used the *Leishmania*-adapted CRISPR/Cas9 system(Zhang and Matlashewski, 2015) to expedite disruption of the AAP3 genes.

Wild type (WT) *L. donovani* was transfected (separately) with plasmids containing three 21-nt gRNAs (G1, G2 and G3) targeting the 5’ end (+19, 85 and 173) of both *AAP3.1* and *AAP3.2* CDSs (Fig. 1A, top panel) and the cultures examined 6 weeks later for AAP3 protein levels in the presence or absence of arginine. While the G1-transfected cultures showed enhanced ADR-mediated increase in AAP3 levels compare to WT, the G2 and G3 cultures showed a diminished capacity to up-regulate AAP3 levels after arginine starvation (Fig. 1B, top left panel). The G3 culture was seeded onto agar plates and individual colonies examined for ADR, with different clones showing normal (*Δap3G3-2*), enhanced (*Δap3G3-17*), or reduced (*Δap3G3-1*) increases in AAP3 protein abundance compared to WT (Fig. 1B, top right panel). In order to increase the efficiency of CRISPR/Cas9-mediated gene disruption, *Δap3G3-1* was transfected (four times at 3-day intervals) with a 61-nt “donor” oligonucleotide consisting of 25-nt flanking an 11-nt insertion with two in-frame stop codons at the Cas9 cleavage site (Fig. 1A, bottom panel). Single colonies were isolated 3 weeks after transfection and examined for AAP3 expression levels after arginine deprivation. Two clones (*Δap3D6* and *Δap3D10*) showed no increase in AAP3 protein abundance after arginine deprivation, while two others (*Δap3D4* and *D7*) showed a small (compared to WT) increase in AAP3 protein levels (Fig. 1B, bottom panel). Analyses of the initial rates of arginine transport confirmed that these mutants had lost all or some of their response to arginine deprivation (Figs. 1C and S1). While WT cells showed a 2.7-fold increase in the initial rate of arginine transport 2 hours after arginine starvation, most of the mutants showed little or no increase in transport rate after arginine starvation (1.5-, 1.2, 1.5- and 1.1-fold for *Δap3D4, Δap3D6, Δap3D7 and Δap3D10*, respectively). Interestingly, *Δap3G3-17* showed a higher than WT (3.2-fold) increase in transport after arginine deprivation, consistent with its having additional copies of the *AAP3.2* gene (see below).

**Figure 1:**
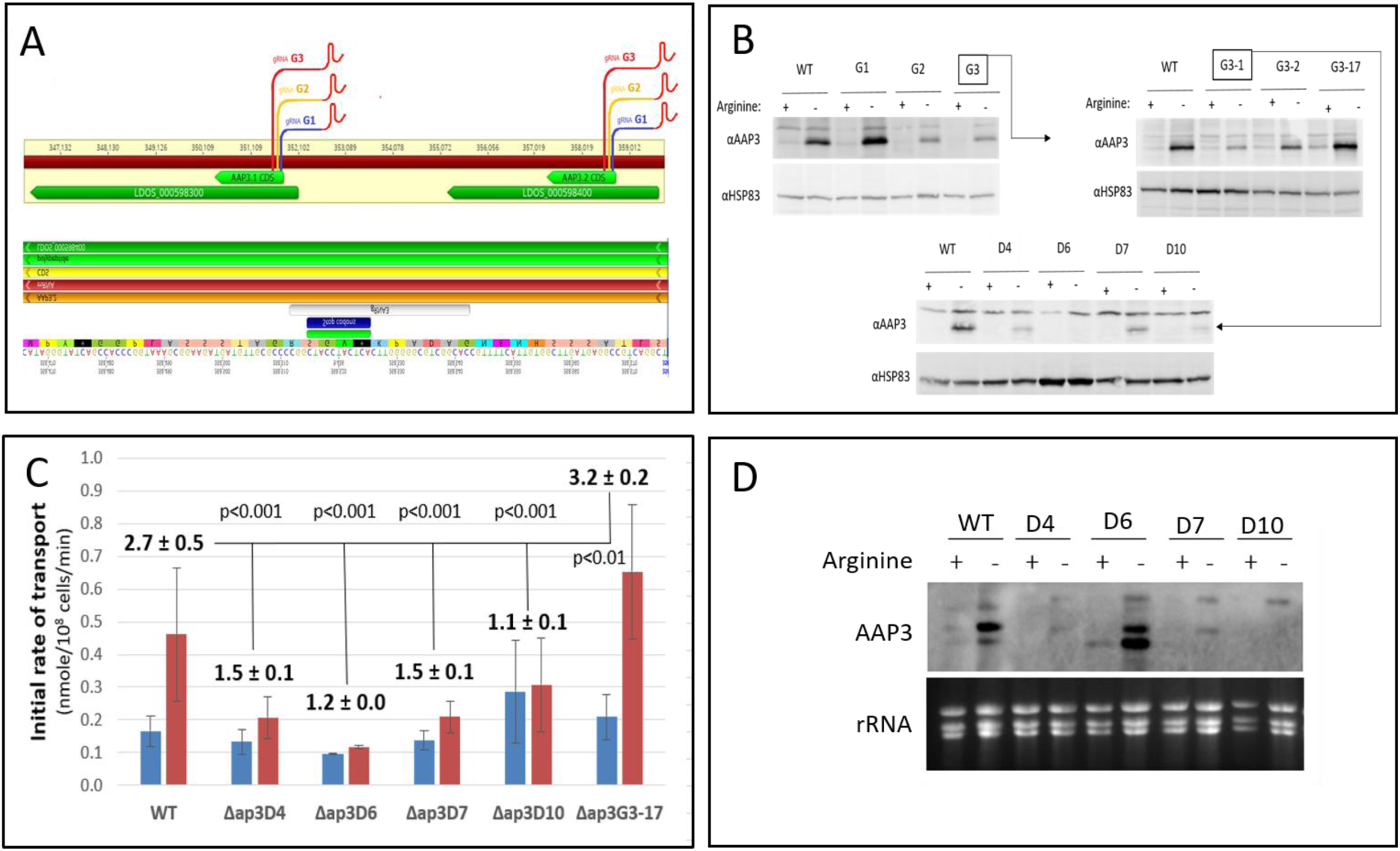
Targeting CRISPR guides to the 5’ end of *AAP3* ORF impaired their response to arginine deprivation. *Leishmania*-adapted CRISPR/Cas9 targeted to the 5’ end of *AAP3* ORF at positions 19, 85 and 173 bp into the CDS with gRNAs designated G1, G2 and G3, respectively (**A, top panel**). gRNA-transfected promastigotes were grown in culture, supplemented with neomycine (50 µg/ml), for six weeks. Subsequently, subjected protein extracted from these cells were subjected to western analysis using anti LdAAP3 antiserum (and HSP83 antisera for loading control, **B, top left panel**). G3 cells (highlighted in square) then seeded on agar plates and three types of colonies raised, *Δap3G3-1*, *Δap3*G3-2 and *Δap3*G3-17 (**B, top right panel**). *Δap3G3-1* (highlighted in square) was then further transfected with a donor, i.e. the gRNA G3 based sequence containing an 11-bp insertion with stop codons (**A, bottom panel**). These transfectants gave rise to several colonies of Δ(*aa3-adr*) mutants that we named *Δap3D4*, *Δap3D6*, *Δap3D7* and *Δap3D10* (**B, bottom panel**). Fold change of initial rate of arginine transport (2 minutes, Fig. S1) before and after two hours after arginine starvation (**C, blue and red, respectively**). (**D**) Northern analysis of AAP3 mRNA expression before and after two hours of arginine starvation.

Whole genome sequencing was carried out to map the precise location of the CRISPR/Cas9-induced mutations in each cell-line. Illumina libraries were constructed and sequenced on a HiSeq 4000 using paired-end 75-bp reads with TruSeq standard primers. Between 45.9M (*Δap3D4*) and 96.3M (*Δap3G3-17*) reads were obtained per library, and they were aligned against an *L. donovani* 1S genome sequence assembled from a combination of PacBio and Illumina reads (manuscript in preparation). As shown in Fig. 2A, changes in coverage (read counts) were seen in the *AAP3* locus of all mutants (*Δap3D7* is not shown since it was essentially identical *Δap3D4*). As expected, the WT parent had four copies of both *AAP3.1* and *AAP3.2*, as well as the genes flanking this locus (Fig. 2B). However, *Δap3G3-1* and most of its progeny (*D4* and *D10*) had substantially lower coverage in the sequence between the 5’ UTR of *AAP3.1* and the 3’ end of *AAP3.2*, consistent with recombination near the double-strand break(s) introduced by Cas9 at the site of the gRNA-G3 sequence to create a *AAP3.1*/*AAP3.2 fusion* that contains the 5’ UTR of *AAP3.*2 and the 3’ UTR of *AAP32*, along with deletion of 3’ UTR of *AAP3.*2and the ncRNA located in the intergenic region between the two *AAP3* genes. Quantitation of the read counts (Fig. 2B) indicated that *Δap3G3-1* retained two intact copies of *AAP3.1* and *AAP3.2*, along with two copies of the *AAP3.1*/*AAP3.2 fusion*; *Δap3G3-1/D4* (and *Δap3G3-1/D7)* retained one copy of the intact *AAP3* locus (containing both *AAP3.1* and *AAP3.2)* and three copies with the *AAP3.1*/*AAP3.2* fusion; while in *Δap3D10* all four copies of the *AAP3* locus contain the *AAP3.1*/*AAP3.2* fusion. However, the presence of a small number (102) of reads from this region in *Δap3D10* suggests that some cells in this population may retain an intact copy of chr31 (although this could be due to presence of an extrachromosomal circle formed between *AAP3.1* and *AAP3.2* or contamination with another cell-line). In contrast, *Δap3G3-17* showed higher coverage in this region, consistent with recombination between the two copies of the *AAP3* gene resulting in additional (2-3 for each copy of chr31) copies of *AAP3.2* and the intergenic sequence (including the ncRNA). Interestingly, *Δap3D6* shows slightly higher than WT levels of read coverage for the *AAP3.1* gene, suggesting that it retained only one copy of chr31 with the *AAP3.1*/*AAP3.2* fusion, along with three copies containing both *AAP3.1* and *AAP3.2* genes (one may contain an additional *AAP3.2* gene). Eexamination of the read coverage in the gRNA-G3 region indicated that ~78% (418/536) contain an 11-bp insertion (with two in-frame stop codons), consistent with the hypothesis that *Δap3D6* contained 7 copies of the *AAP3* gene with stop codons that render them non-functional at the protein level and only one (or two) functional version(s). The inability of this mutant to present with even partial ADR on protein level such as that of mutants D4/D7 suggests that the functional copies are the fused *AAP3.1*/*AAP3.2* which lack stop codons but also lack ADR capacity.

**Figure 2:**
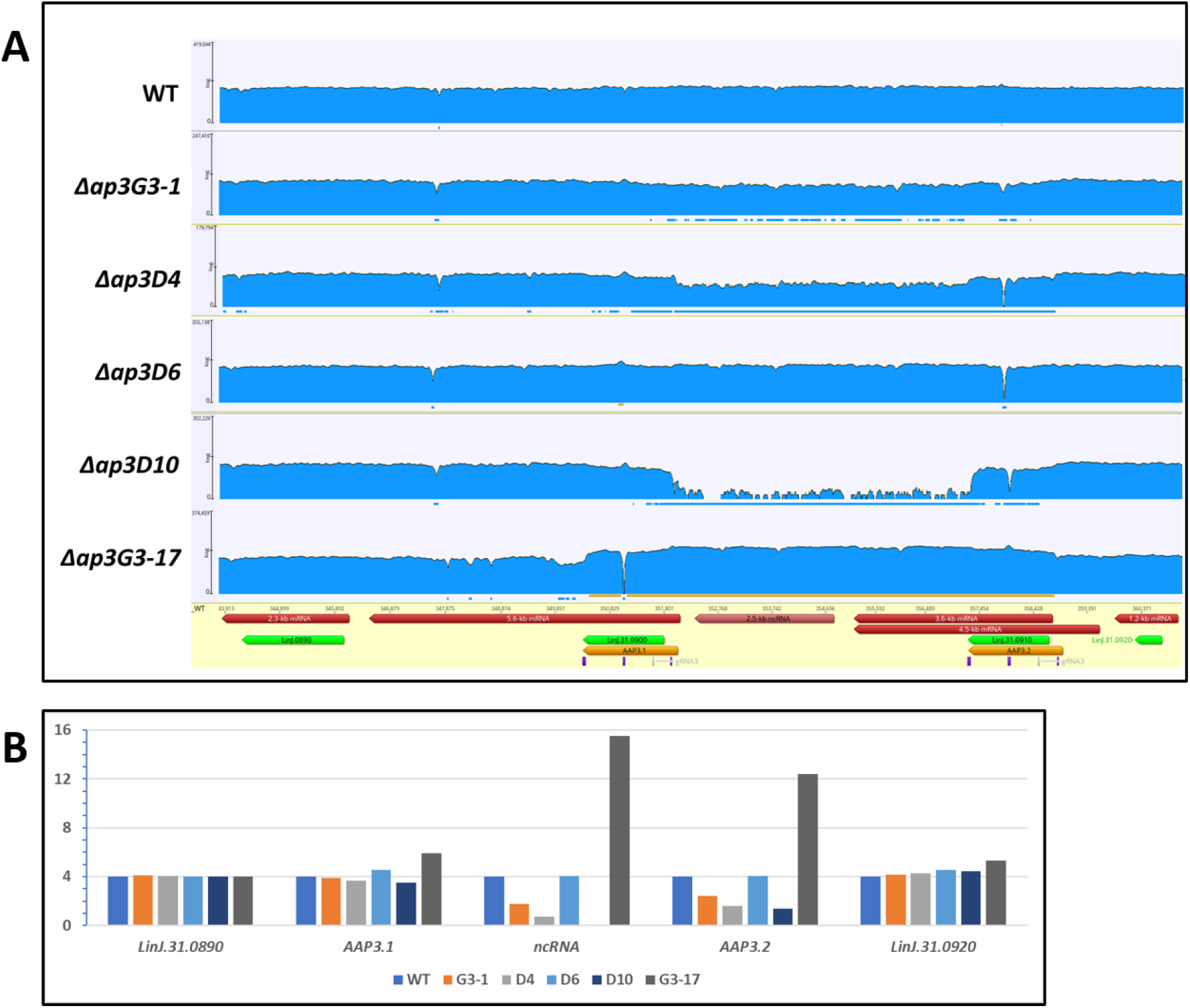
Whole genome sequencing of the AAP3 region in the CRISPR/Cas9 treated L. donovani promastigotes. Whole genome sequencing carried out for each of the WT and *Δ(aap3-adr)* mutants. Illumina libraries were constructed and sequenced on a HiSeq 4000 using paired-end 75-bp reads with TruSeq standard primers. (**A**) Reads were aligned against the Ldo1S reference genome and revealed that mutants *Δap3G3-1*, *Δap3D4* and *Δap3D10* harbor a large deletion essentially fusing AAP3.1 CDS with that of AAP3.2 and eliminating AAP3.2 3’UTR and all intergenic ncRNA sequences. Quantification of reads indicates that gene fusion event has occurred on two copies of chromosome 31 in *Δap3G3-1*, three copies in *Δap3D4* and on four copies in *Δap3D10*. *Δap3G3-17* harbors 16 copies of the region. *Δap3D6* alignment to a version of LdoS_31 revealed an 11-bp insertion (with a stop codon in each frame) in the AAP3.2 gene. The read counts for each gene represent only those that can be unambiguously assigned to the WT or mutant version of the gene. These results are consistent with the hypothesis that 3 copies of AAP3.2 and 3 copies of the 4 AAP3.1 genes contain stop codons in *Δap3D6* (**A**). Reads analysis indicated that neighboring genes such as LinJ.31.0890 and LinJ. 0920 were not affected by the AAP3 mutation (**B**).

Northern analyses (Fig. 1D) confirmed the reduction in *AAP3.2* gene copy number by showing that its mRNA levels after arginine deprivation were significantly lower than WT in the in *D4*, *D7* and *D10* cell-lines. These changes paralleled the changes in protein abundance observed in Fig. 1B, confirming that AAP3.2 accounts for most (if not all) of the increase in arginine transport after arginine starvation. Importantly, even though the *AAP3.2* mRNA levels in *Δap3D6* were similar to WT levels in both the presence and absence of arginine, AAP3 protein levels were not up-regulated in response to arginine deprivation because of the stop codons in the *AAP3.2* gene. Two other ADR-responsive genes(Goldman-Pinkovich et al., 2016), *LinJ.10.1450* and *LinJ.36.2900* (which encode pteridine and MSF family transporters, respectively), responded normally to arginine deprivation in all mutants (Fig. S2), indicating that the CRISPR/Cas9 mutations affected only the response of *AAP3.2* to arginine deprivation, not the entire ADR pathway.

The analyses described above indicated that *Δap3D6* and *Δap3D10* are the most informative *AAP3* mutants, having retained near-WT basal levels of arginine transport (when grown in normal medium), but being unable to up-regulate transporter activity after arginine deprivation. Therefore, we decided to pursue virulence studies with these two mutants to assess whether increased AAP3 expression is essential for infectivity and virulence. We infected THP1 macrophages with late-log phase *L. donovani* promastigotes of WT, *Δap3D6, Δap3D10* and *Δap3G3-17* in medium containing a physiologically-relevant concentration of arginine (0.1 mM) (Brown et al.),(Pawar et al., 2019). As shown in Fig. 3A, the initial (after 4 hours of co-incubation) level of infection (*i.e.* the percentage of macrophages infected, upper panel) and parasitemia (*i.e.* the average number of parasites per macrophage, lower panel) with all three mutants were similar to those of WT. However, by 48 hours post-infection, the infectivity and parasitemia for both *Δap3D6* and *Δap3D10* were reduced by 2-fold or more compared to WT, while the levels in *Δap3*G3-17 were comparable to WT (Fig. 3B). Hence, our results indicate that the mutants are able to infect macrophages (almost) as well as WT, they fail to proliferate normally thereafter. Indeed, it appears that significant number of macrophages are able to completely clear their internalized parasites, while the parasites in the remaining infected macrophages undergo only one or two rounds of replication (compared to the normal 3-4 rounds of replication and re-infection of new macrophages seen with WT parasites).

**Figure 3:**
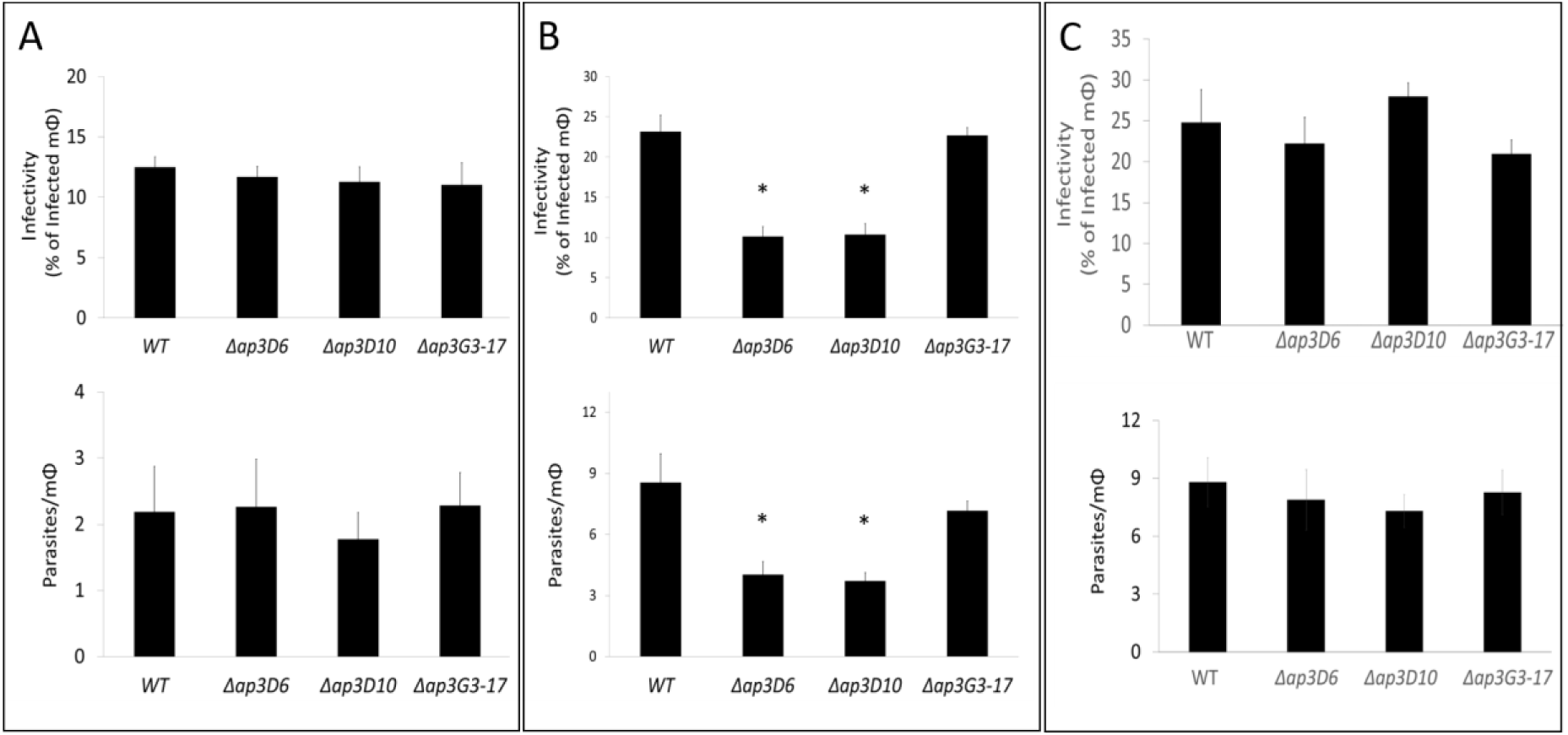
Mutants lacking AAP3-ADR lost infectivity to THP-1 macrophages. Human THP-1 differentiated macrophages grown in RPMI1640 medium containing 0.1 mM arginine, subjected to infection by mid-log *L. donovani* promastigotes of WT, *Δap3D6*, *Δap3D10* and *Δap3G3-17* at an MOI of 10 for four hours, as described in Methods. At times zero and 48 hours after infection macrophages were fixed, giemsa stained and subsequently subjected to counting. Virulence capacity was determined by calculating infectivity (A and B top panels) and parasitemia (A and B bottom panels). Macrophage infections carried out in the presence of 0.1 mM arginine (A and B) or 1.5 mM (C). For panel B, one-way ANOVA indicated that *Δap3D10* and *Δap3D6* mutants were 57% less infective than WT (p<0.05, n=16) and parasitemia 50% less than WT (p<0.05, n=4). When infection was performed at 1.5 mM external arginine concentration One-way ANOVA was insignificant for mutant impairment in either infectivity (**C, top panel**) or parasitic burden (**C, bottom panel**) parameters (p<0.1).

As we have previously shown (Pawar et al., 2019), 24 hours after infection of macrophages grown in a medium containing 0.1 mM arginine, amastigotes activate the ADR likely due to arginine depletion in the phagolysosome of infected macrophages. Thus, we used qRT-PCR to quantify mRNA levels in promastigotes immediately before infection of THP1 cells and amastigotes 48h after infection. As expected, WT amastigotes showed a significant (~3-fold) increase in *AAP3.2* mRNA levels when compared to promastigotes, but the *AAP3.2* mRNA levels in amastigotes from the *Δap3D6* and *Δap3D10* mutants were substantially lower than WT. In addition, it should be remembered that the *AAP3.2* mRNA in the former is non-functional due to in-frame stop codons. In contrast, *AAP3.2* mRNA levels in the *Δap3*G3-17 mutants (even promastigotes) were higher than WT amastigotes. To assess whether the lower AAP3 expression levels (and consequently less arginine import) was responsible for the reduction in infectivity and parasitemia of the *Δap3D6* and *Δap3D10* mutants, we infected THP1 macrophages grown in medium containing 1.5 mM arginine, a concentration that prevented ADR in intracellular WT amastigotes (Pawar et al., 2019). Under these conditions, both *Δap3D6 and Δap3D10* developed normally into amastigotes, with similar infectivity and parasitemia at 48 hours to that of WT (Fig. 3C). The results indicate that the inability to develop inside macrophages is due to arginine deprivation that developed in macrophage phagolysosomes during infection and that the ability to respond to this deprivation by up-regulating *AAP3.2* protein levels is essential for successful intracellular *Leishmania* development.

To further assess the role of arginine transport *in vivo*, we infected BALB/c mice with *Δap3D6, Δap3D10*, *Δap3G3-17* and WT (8 mice for each group). On day 21 post-infection, the mice were sacrificed and parasitemia determined by qPCR on the DNA extracted from the liver of each mouse. As shown in Fig. 4, the parasitemia of the *Δap3D6* and *Δap3D10* mutants were only 20 and 24%, respectively, that of WT. One-way ANOVA and Tukey post-hoc HSD testing showed a significant difference (P<0.001) from WT for both the *Δap3D6* and *Δap3D10* mutants, while *Δap3G3-17* was not significantly different from WT (p=0.387). These results show that inability of the mutants to express higher levels of AAP3 in order to compensate for the reduced level of arginine in the phagolysomes of infected macrophages severely compromised their virulence. Therefore, it appears that the ADR is a crucial mechanism for enabling intracellular *Leishmania* parasites to overcome the arginine bottleneck and win the “Hunger Games” in the mammalian host.

**Figure 4.**
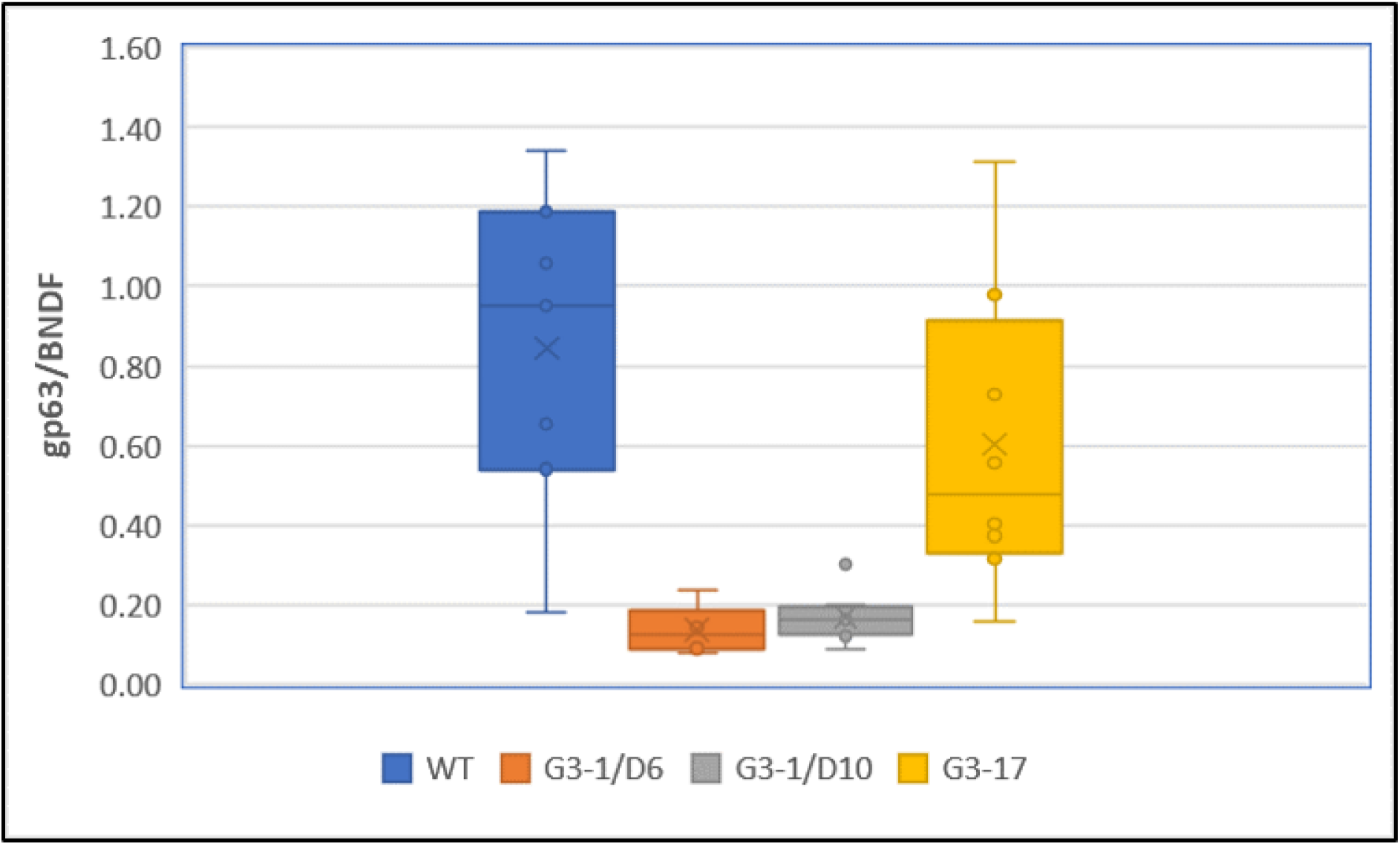
*AAP3.2* mutants are unable to develop in mice. Late stationary phase *L*. *donovani* promastigotes of WT, *Δap3D6*, *Δap3D10* and *Δap3G3-17* injected intravenously (IV) into eight-week-old female BALB/c mice (1×10^8^ cells per injection, eight mice per mutant). On day21 post infection, mice were sacrificed and liver DNA extracted using the proteinase K method(Hofstetter et al., 1997). Analyses were carried out as described in Methods.

In summary, this study demonstrates for the first time that the ability to monitor metabolic deprivation and subsequently induce a specific response at the level of gene expression is essential for pathogenesis of a protozoan parasite. Furthermore, we have shown that in order for pathogenic microorganisms to respond to host-inflicted environmental changes and survive, they must employ external sensing and response mechanisms that serve as their monitoring device.

## Materials and Methods

### Leishmania strains and culture

*L. donovani* MHOM/SD/00/1S (Saar et al., 1998) promastigotes were grown in Medium M-199, Earle’s salts (Biological Industries Ltd) supplemented with 10% heat-inactivated Fetal Bovine Serum (GIBCO Ltd) and 1% Penicillin-Streptomycin solution (Biological Industries Ltd). Arginine deprivation was carried out as described by Pawar et al. (Pawar et al., 2019). Briefly, mid-log phase promastigotes (1×10^7^ cells/ml) were washed with Earle’s balanced salt solution twice and re-suspended in arginine-deficient Medium M-199 (Biological Industries Ltd.) at 26°C for the specified period before being transferring to ice. Arginine-deprived cells were washed twice with ice-cold Earle’s balanced salt solution before being used for transport assays, Northern and Western blot analysis.

### CRISPR/Cas9 guide and donor transfections

gRNA sequences were designed using the Eukaryotic Pathogen CRISPR guide RNA/DNA Design Tool (EuPaGDT, http://grna.ctegd.uga.edu/) and cloned into *Leishmania*-adapted vector pLdCN using the single-step digestion-ligation cloning protocol previously described (Zhang and Matlashewski, 2015), and the constructs transfected into mid-log phase promastigotes. Following gRNA-pLdCN transfections, cells were grown for 4 weeks with G418 at 50 μg/ml and subsequently screened for their ability to increase LdAAP3 protein abundance after arginine deprivation. Next, the G3-donor sequence was introduced into a gRNA G3-originated clone exhibiting the desired phenotype. The G3-donor is a single strand oligo donor (sense) containing 25-nt sequences flanking the Cas9 cleavage site (shown in bold below) and an 11-nt sequence with two in-frame stop codons (underlined and italicized below). Three transfections with 10 µl (100 µm) of this oligo were performed on promastigotes at three day intervals as previously described (Zhang and Matlashewski, 2015). gRNA sequences.

gRNA G1: GTCTATTCCAGCACAGGCGG

gRNA G2: GCCGTCGATAAACACCCGAG

gRNA G3: GTGCCGACGCCGCCAAGCCG

G3-based donor:

ATGAAAACG**GTGCCGACGCCGCCAAG**<u>*TGAGTAGGTAG*</u>**CCG**GGGCGCAACATCATCTTCCG

### Western Blot Analyses

Western blot analysis was done as described previously(Darlyuk et al., 2009) using a 1:1000 dilution of rabbit anti AAP3 N-terminus antisera(Goldman-Pinkovich et al., 2016).

### Transport assays

Uptake of 25 μM L-[^3^H] arginine (600 mCi/mmol) into *L. donovani* promastigotes was determined using the rapid filtration technique described previously(Goldman-Pinkovich et al., 2016). Briefly, transport reaction mixture contained 1×10^8^ cells/ml suspended in Earle’s balanced salt solution at pH 7 supplemented with 5 mM glucose and 1% dialyzed heat-inactivated fetal calf. Cells were pre-warmed at 30°C for 10 min prior to addition of radiolabeled arginine. After 0, 0.5, 1, 1.5 and 2 minutes, the cell suspensions were filtered through GF/C filters. The amount of radiolabel associated with the cells was linear with time over the 2-min time course of the assay, so the initial rate of transport was calculated from the slope of the line fitted by linear regression (Fig. S1).

### RNA isolation and real time quantitative reverse transcription PCR

RNA was isolated using TRI reagent® (Sigma-Aldrich Ltd) or Direct-zol RNA MiniPrep Kit (Zymo Research), according to manufacturer instructions. Eluted RNA samples were quantified using a NanoDrop One spectrophotometer (Thermo Scientific). Two μg of the extracted RNA were subjected to DNase treatment using RQ1 (Promega). Successful DNase treatment was verified by PCR to make sure no residual DNA could be responsible for amplification. cDNA was synthesized from 2 μg of DNase-treated RNA using a qScript cDNA Synthesis Kit (Quanta Biosciences) in 40 μl total volume. qRT-PCR was carried out with the reagents of SsoAdvanced Universal SYBR Green Supermix (Bio-Rad, Ltd.) in 10 μl reaction volume (5 μl SYBR Green, 0.5 nM forward primers, 0.5 nM reverse primers and 2.5 μl cDNA template) on CFX96 Touch Real-Time PCR system (Bio-Rad). The AAP3 primers matched both *AAP3.1* and *AAP3.2*. Primers specific for a regulatory subunit of PKA (PKAR′) were used as a control. All the samples were run in triplicates, including a no-template (negative) control for all primers used. Also, RNA-seq data of Arginine Deprivation Response indicated PKAR’ is not affected by ADR thus serving as a good control for testing LdAAP3 behavior under ADR and related conditions.

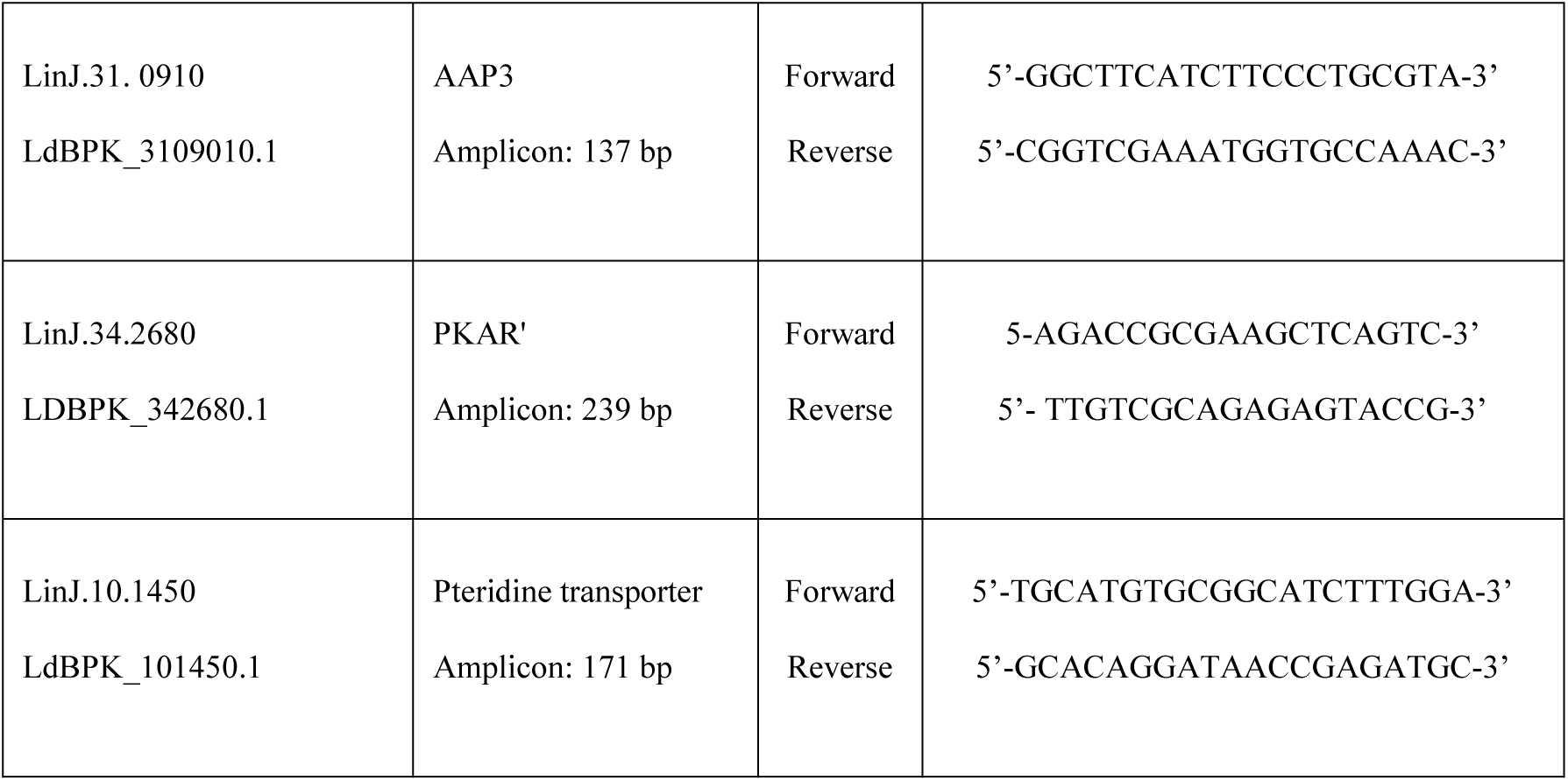
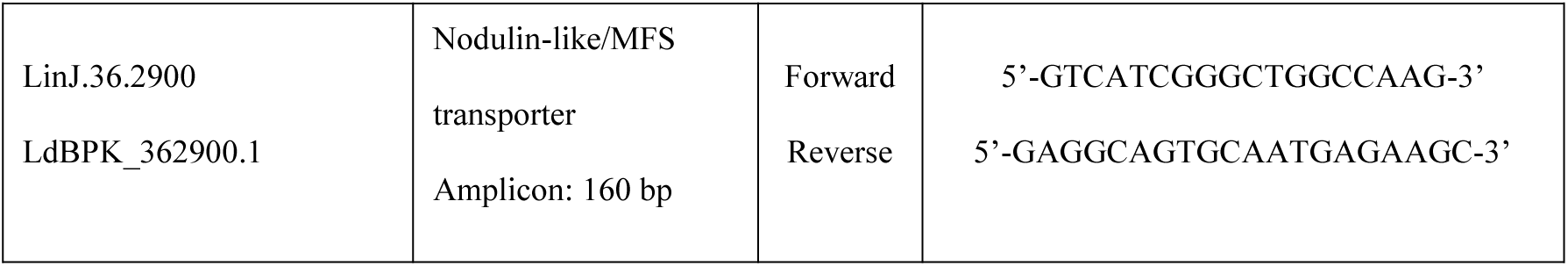

PCR was performed at 95°C for 30 seconds, followed by 39 cycles of 95°C for 10 seconds, 60°C for 30 seconds and Melt Curve analysis was carried out at 65-95°C with 0.5°C increment for 5 sec/step and data analysis was carried out as described previously(Pfaffl, 2001). Briefly, C_P_ values were obtained from all samples in triplicate (except negative controls). Primers were calibrated on pooled samples and primer efficiencies (E_target_ and E_ref_) were calculated and incorporated in the equation below where LdAAP3, Pteridine or MFS transporters were the target gene and PKAR’ served as the reference gene. Ratios were calculated, with C_P_ of 0 hours or 48 hours infected THP-1 macrophages. WT *L. donovani* 48 h infected macrophages served as control.

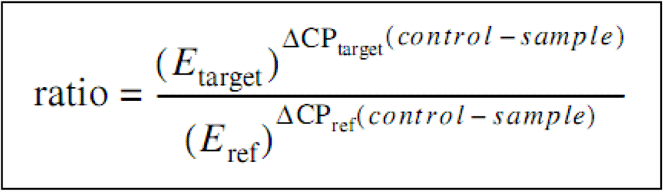

### Whole Genome Sequencing

Genomic DNA was prepared from *Leishmania* promastigotes using proteinase K digestion followed by phenol/chloroform extraction and ethanol precipitation and fragmentation to 200bp using a Covaris S2 sonicator according to the manufacturer’s protocol. NGS libraries were prepared using an NEB NextUltra II kit, quantified on Agilent Bioanalyzer and Qubit instruments and sequenced on an Illumina Hiseq 4000 to generate 46-96 million paired-end 75-base pair reads, depending on the sample. Reads were aligned to the *L. donovani* 1S genome sequence assembled from a combination of PacBio and Illumina reads (manuscript in preparation) using Bowtie2 within Geneious and gene copy number estimated by normalizing the read counts per gene to account for library size and assuming 4 copies of chr31.

### Infectivity assays

THP-1 cells were seeded on glass coverslips (1×10^6^ cells/well) in a 6-well plate and treated with 50 ng/ml of PMA (Sigma-Aldrich, USA) for 48 h. They were infected as described previously (Pawar et al., 2019) and the intracellular parasite load was visualized by Giemsa staining.

### In vivo BALB/c mice infections and analysis

L. *donovani* infections were done by tail intravenous (IV) injection of 10^8^ stationary phase promastigotes per mouse. On day 21 of the infection the mice were sacrificed and mouse liver DNA isolated using the protein K method (Hofstetter et al., 1997). Quantitative PCR was performed on mouse liver DNA using mouse BDNF as reference genes and parasite gp63 as target gene with published primer sequences (Tiwananthagorn et al., 2012),(Tupperwar et al., 2008).

Q-PCR was carried out with the reagents of SYBR Green (ThermoFisher Scientific, Catalog No A25776) in 10 μl reaction volume (5 μl SYBR Green, primer final concentration of 300 nM forward primer, 300 nM reverse primer and 125ng DNA template) on CFX96 Real-Time PCR system (Bio-Rad). All samples were run in triplicate separately for the 2 primer sets and the PCR cycle include 30 s incubation step at 95°C, then 40 cycles of 5 s at 95 °C and 30 s at 60 °C. The output of normalized expression was determined by the BIORAD software of the instrument.

## Acknowledgments

We thank Prof. Ido Izhaki of Haifa University for his help with the statistical analyses and MS Michal Almoznino for experimental analyses.

Funding for this work was provided by the University Grant Commission (UGC) – Israel Science Foundation (Indo-Israel) research grant number 2316/15 to DZ, AG and RM and USA-Israel Binational Foundation grant number 2017030 to DZ and PJM.

---

**Figure S1:**
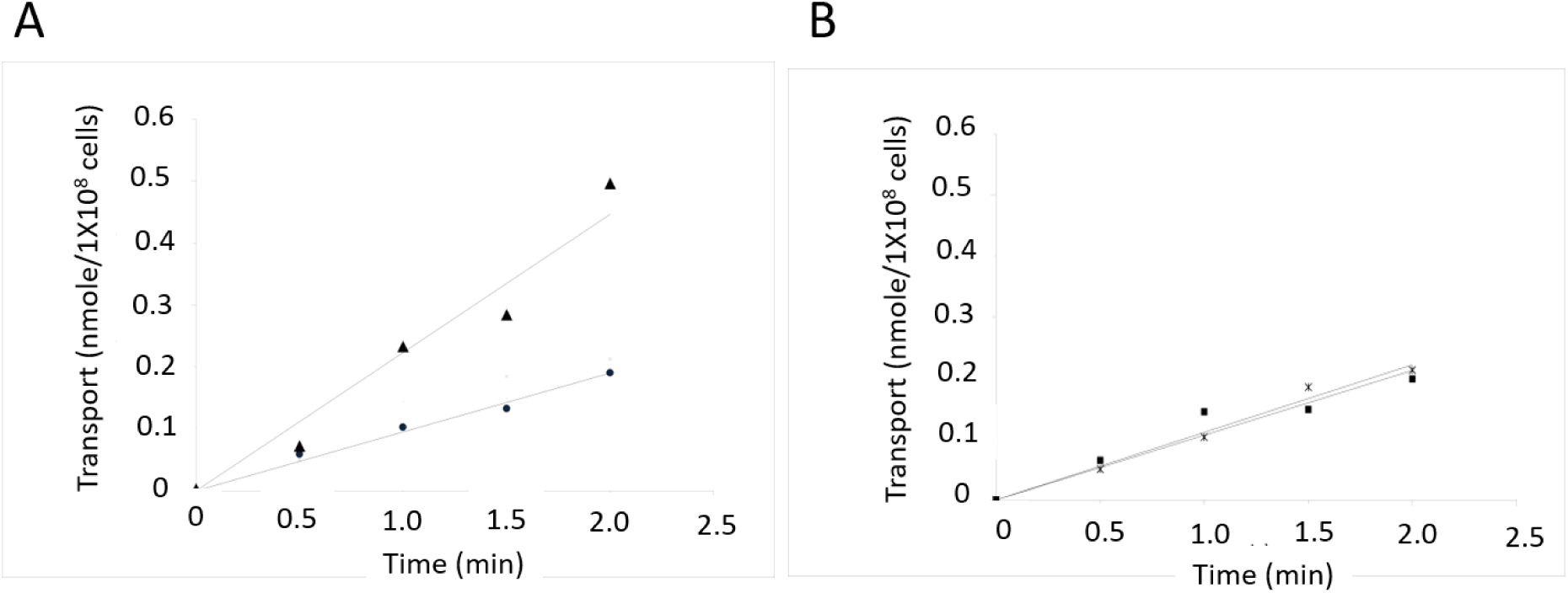
Initial rate of arginine transport in wild type and mutant *L. donovani* promastigotes. Two minutes arginine transport was determined using the rapid filtration technique (Methods). Transport was assayed in wild type before (●) and after (▲) two hours arginine starvation and in *Δap3D6* before (●) and two hours after arginine starvation 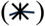.

**Figure S2:**
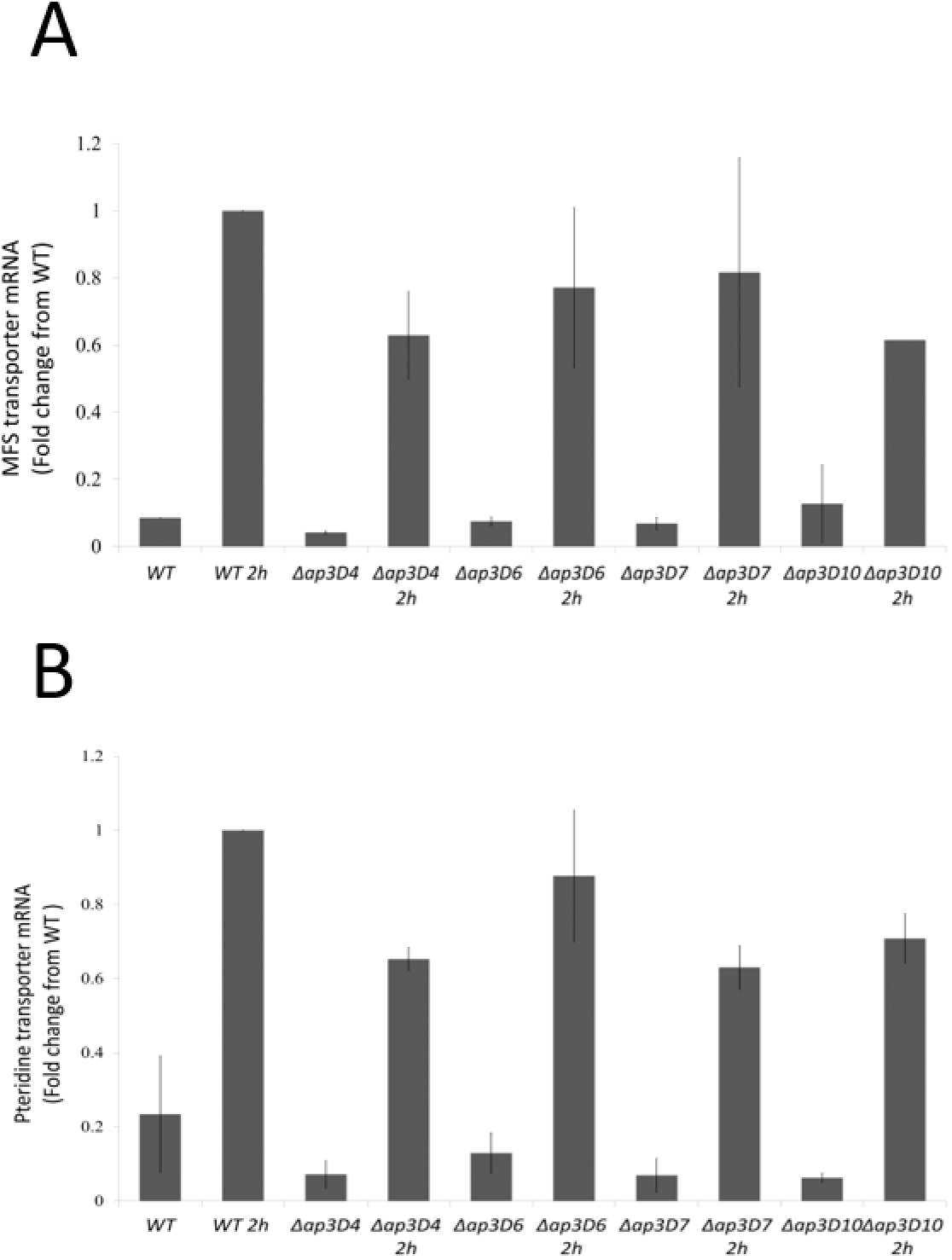
Other members of ADR retained sensitivity to arginine deprivation. Real Time PCR analysis was carried out to determine mRNA levels of pteridine (upper panel) and MSF (lower panel) transporters before and after two hours after arginine starvation. The analyses were carried out in *Δap3D4, Δap3D6, Δap3D7 and Δap3D10.* Results are relative to 2 hour arginine deprived WT cells.

